# P-bodies are sites of rapid RNA decay during the neural crest epithelial—mesenchymal transition

**DOI:** 10.1101/2020.07.31.231860

**Authors:** Erica J. Hutchins, Michael L. Piacentino, Marianne E. Bronner

## Abstract

The epithelial—mesenchymal transition (EMT) drives cellular movements during development to create specialized tissues and structures in metazoans, using mechanisms often coopted during metastasis. Neural crest cells are a multipotent stem cell population that undergo a developmentally regulated EMT and are prone to metastasis in the adult, providing an excellent model to study cell state changes and mechanisms underlying EMT. A hallmark of neural crest EMT during avian development is temporally restricted expression followed by rapid down-regulation of the Wnt antagonist *Draxin*. Using live RNA imaging, here we demonstrate that rapid clearance of *Draxin* transcripts is mediated post-transcriptionally via localization to processing bodies (P-bodies), small cytoplasmic granules which are established sites of RNA processing. Contrasting with recent work in immortalized cell lines suggesting that P-bodies are sites of storage rather than degradation, we show that targeted decay of *Draxin* occurs within P-bodies during neural crest migration. Furthermore, P-body disruption via *DDX6* knockdown inhibits not only endogenous *Draxin* down-regulation but also neural crest EMT *in vivo*. Together, our data highlight a novel and important role for P-bodies in an intact organismal context—controlling a developmental EMT program via post-transcriptional target degradation.

## INTRODUCTION

The epithelial—mesenchymal transition (EMT) is an impactful cell behavior in normal and disease states in metazoans (Yang et al., 2020). Cell movements that are a product of EMT during embryonic development are essential to form numerous tissues and organs via mechanisms that are analogous to cell invasion during cancer metastasis (Acloque et al., 2009; Hutchins et al., 2018; Kerosuo and Bronner-Fraser, 2012; Theveneau and Mayor, 2012; Thiery et al., 2009). The EMT program is dynamic and requires rapid molecular transitions to achieve a spectrum of intermediate cell states that imbue cells with different degrees of plasticity, reversibility, invasion, and migration (Aiello et al., 2018; Nieto et al., 2016; Pastushenko and Blanpain, 2019). The neural crest is a classic example of an essential developmental cell type that undergoes a tightly regulated EMT driven by changing molecular signatures that alter their migratory ability, and ultimately cell fate choice (Gouignard et al., 2018; Martik and Bronner, 2017). These intrinsic cellular changes are driven by sequential activation of gene regulatory networks (Fazilaty et al., 2019; Williams et al., 2019), the outputs of which are subsequently fine-tuned via post-transcriptional regulatory mechanisms (Bhattacharya et al., 2018; Weiner, 2018).

In both development and disease, post-transcriptional regulation and, in particular, RNA turnover are key mechanisms by which cells are able to quickly alter their transcriptomic landscape and transit between cell states (Chen and Shyu, 2017; Hardy et al., 2017; Lou et al., 2015). The process of RNA decay is highly conserved across eukaryotes and mediated by multiple mRNA turnover pathways (Parker and Song, 2004). One of the major eukaryotic mRNA decay pathways is Xrn1-mediated, which occurs 5’→3’ as opposed to 3’→5’ exosome-mediated decay (Muhlrad et al., 1995; Siwaszek et al., 2014). Xrn1-mediated decay was originally demonstrated in yeast to occur within discrete cytoplasmic foci called processing bodies (P-bodies) (Sheth and Parker, 2003). Interestingly, P-body formation is necessary for EMT in cancer cells; disrupting P-bodies blocked metastasis and caused the cells to retain more epithelial characteristics (Hardy et al., 2017). However, which transcripts localize to P-bodies during EMT or how they are processed to drive mesenchymal characteristics remain unknown. Recent work in human cell lines has suggested that metazoan Xrn1-mediated decay may occur broadly within the cytosol, with P-bodies serving as sites of translational repression and storage rather than decay (Horvathova et al., 2017; Hubstenberger et al., 2017), though this has been shown to vary based on target and context (Aizer et al., 2014). Thus, the role of P-bodies during EMT as sites of storage versus decay remains controversial.

Here, we describe a critical role for P-body-mediated RNA decay in control of an essential vertebrate developmental EMT program—onset of neural crest cell migration. Using adapted live RNA imaging tools, we show that transcripts encoding Draxin, a molecular rheostat controlling neural crest EMT, are targeted to and decayed within P-bodies in neural crest cells via a DDX6-dependent mechanism.

## RESULTS

### An EMT rheostat is regulated via RNA decay

Cranial neural crest EMT in the chick embryo is spatiotemporally controlled by transient attenuation of canonical Wnt signaling (Hutchins and Bronner, 2018, 2019; Maj et al., 2016; Rabadán et al., 2016). A major regulator of this damping of Wnt signaling is the secreted protein, Draxin, which is expressed in a brief pulse to modulate gene expression output for proper timing of cranial neural crest EMT. A hallmark of *Draxin* function in this process is the rapid down-regulation of its mRNA, coincident with onset of the EMT program (**Fig. 1A-B**); this clearance of transcript is essential for cranial neural crest cells to achieve a mesenchymalized state during EMT (Hutchins and Bronner, 2018, 2019). Using *in situ* hybridization chain reaction (HCR), a highly sensitive method that allows visualization of single mRNA transcripts with subcellular localization, we examined *Draxin* mRNA following cranial neural crest EMT. The results reveal low levels of *Draxin* mRNA localized to discrete cytoplasmic granules, which did not colocalize with *AP2β* mRNA, a transcript highly expressed in migratory neural crest (Simoes-Costa and Bronner, 2016) (**Fig. 1C-D**). The granular appearance of *Draxin* mRNA coincident with its rapid downregulation led to the intriguing hypothesis that *Draxin* may be targeted to cytoplasmic granules for decay (Anderson and Kedersha, 2009) to facilitate EMT.

**Figure 1.**
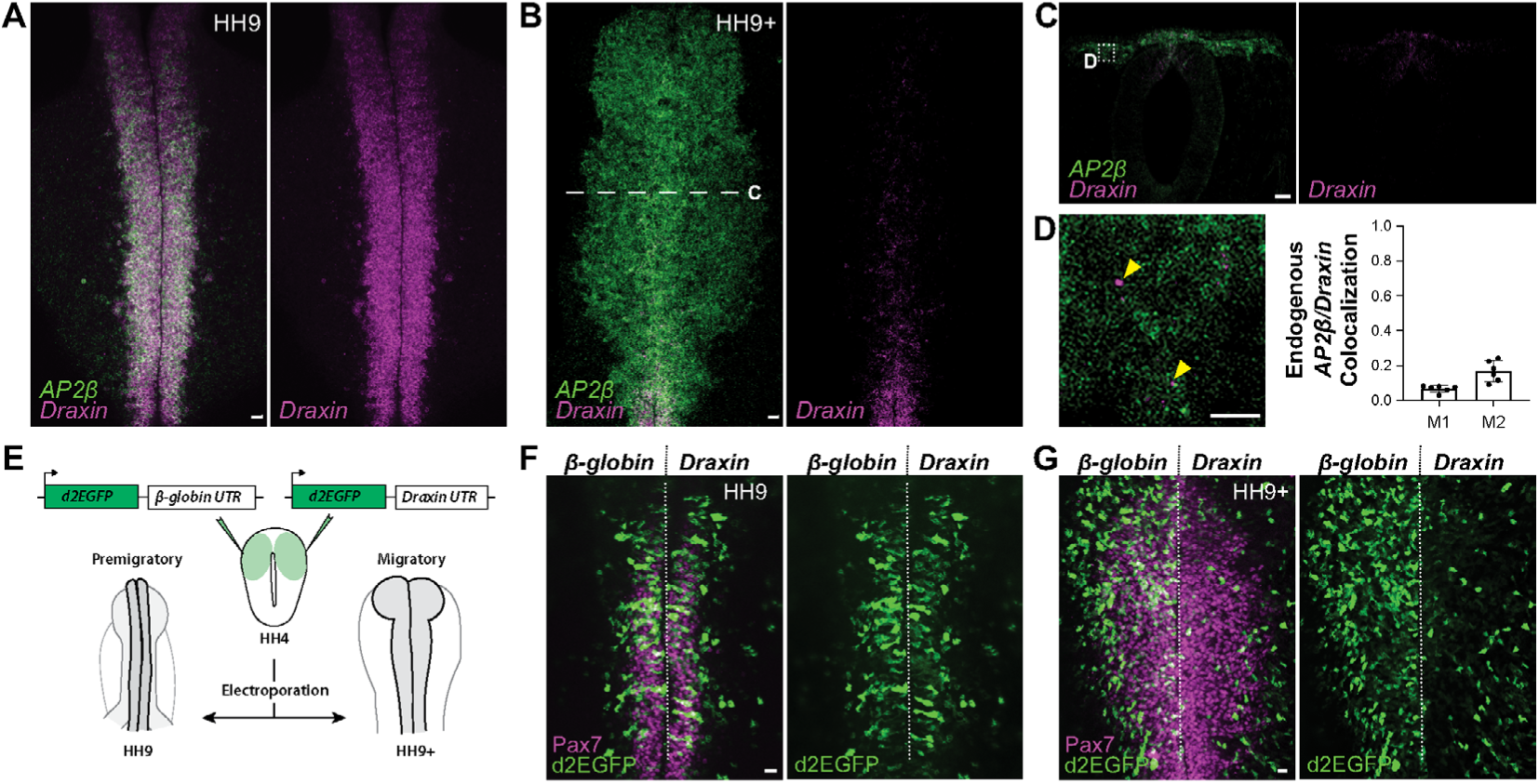
The EMT rheostat *Draxin* is post-transcriptionally regulated. (**A-D**), Representative confocal maximum intensity projection micrographs of HCR processed wild type embryos for *Draxin* and *AP2β* transcripts in whole mount (**A-B**) and section (**C-D**). Mander’s coefficients (M1 and M2) were calculated to determine overlap of expression between *Draxin* and *AP2β* in migratory neural crest. (**E**) Experimental design to examine *Draxin*’s post-transcriptional regulation using UTR-reporter constructs. (**F-G**) Representative epifluorescence images of embryos expressing UTR-reporter constructs and immunostained for neural crest marker Pax7. HH, Hamburger-Hamilton stage; M, Mander’s coefficient; UTR, untranslated region. Scale bars, 20 μm (**A-C,F-G**) and 1 μm (**D**). Error bars, SEM.

To explore this possibility, we next investigated whether the localization and down-regulation of *Draxin* accompanying EMT is mediated by cellular RNA decay pathways leading to targeted degradation of transcript. To this end, we designed an *in vivo* fluorescent reporter construct to first determine whether *Draxin* is post-transcriptionally regulated via its 3’-untranslated region (UTR) within living embryos. The 3’-UTR is a major determinant of mRNA stability, owing to *cis* regulatory sequences (e.g. AU-rich elements) contained therein (Garneau et al., 2007; Grzybowska et al., 2001; Guhaniyogi and Brewer, 2001). We electroporated gastrula stage chick embryos (Hamburger-Hamilton stage HH4 (Hamburger and Hamilton, 1951)) with reporter constructs driving ubiquitous expression of destabilized EGFP (d2EGFP), followed by either a control 3’-UTR (*β-globin*), which generates a stable and highly translated transcript (Lodish and Small, 1976; Wilson and Deeley, 1995), or the endogenous *Draxin* 3’-UTR (**Fig. 1E**). The results revealed d2EGFP expression in premigratory neural crest cells from both constructs. Interestingly, at the migratory stage, only the control 3’-UTR construct displayed robust d2EGFP expression; in contrast, expression from the *Draxin* 3’-UTR construct was drastically diminished, consistent with the timing of endogenous *Draxin* down-regulation (**Fig. 1F-G**). Together, these data suggest a post-transcriptional regulatory mechanism for *Draxin* down-regulation during cranial neural crest EMT, mediated via its 3’-UTR.

### *Draxin* targets to P-bodies during EMT

To understand the dynamics of *Draxin*’s post-transcriptional regulation during EMT in living cells, we adapted the improved, degradable MS2-MCP reporter system (Tutucci et al., 2018) to visualize the control (*β-globin*) and *Draxin* 3’-UTR-containing RNAs by time-lapse imaging. We electroporated early gastrula stage embryos with YFP-tagged, nuclear MS2 coat protein (MCP) in conjunction with a modified MS2 stem loop (24xMBSV6) construct containing either the *β-globin* or *Draxin* 3’-UTR, thus allowing us to infer subcellular localization and stability of transcript based on fluorescence signal intensity from MCP within the cytoplasm. At neurula stages, the cranial neural folds were explanted to enable visualization within emigrating and migrating neural crest cells (**Fig. 2A**). As expected for the control UTR-containing construct (MS2-*β-globin*-UTR), MCP-bound transcripts formed large (≥ 2μm diameter) cytoplasmic granules that were intensely bright and stably fluorescent during cranial neural crest cell migration (**Fig. 2B-C**). Importantly, the *Draxin* 3’-UTR-containing construct (MS2-*Draxin*-UTR) in contrast localized to small (< 1μm diameter) cytoplasmic granules that degraded within 2 h (**Fig. 2B-C**; *P* < 0.001, Kolmogorov-Smirnov nonparametric test). Treatment with the translation inhibitor cycloheximide, which stabilizes ribosomes bound to RNA and dissolves P-bodies (Aizer et al., 2008), had no effect on the large control granules, suggesting these were transcripts associated with ribosomes and likely sites of active translation (**Supplementary Fig. S1A**); notably, however, we saw a decrease in the number of MS2-Draxin-UTR granules (**Supplementary Fig. S1B**), suggesting these transcripts may be localizing to P-bodies for decay during neural crest EMT.

**Figure 2.**
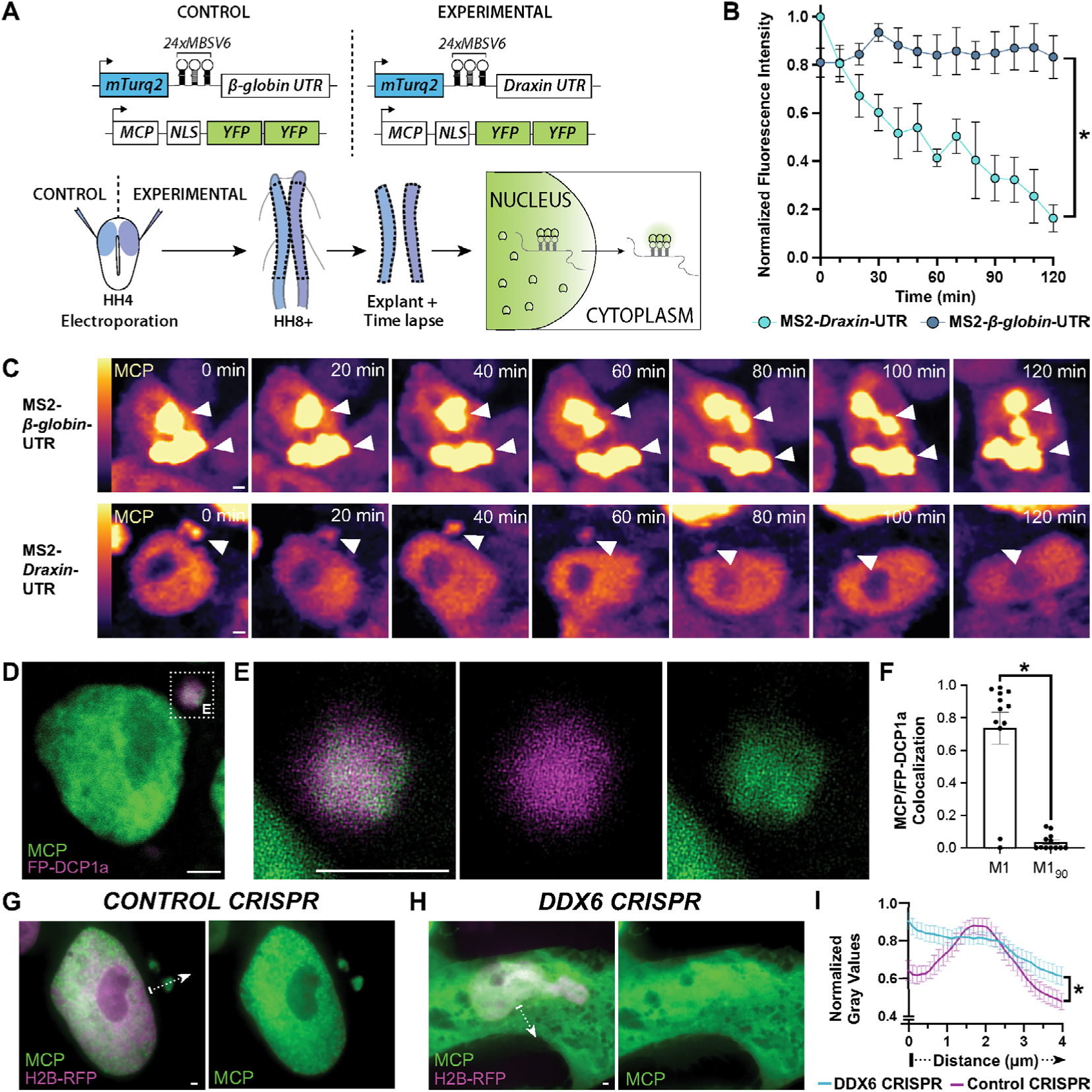
*Draxin* localizes to P-bodies during neural crest EMT. (**A**) Experimental design to visualize transcript dynamics in migrating neural crest cells. (**B-C**) Normalized fluorescence intensity measurements (**B**) of MCP-labeled cytoplasmic granules, determined from confocal time lapse (**C**) of neural crest cells explanted from embryos co-electroporated with YFP-tagged, nuclear MCP and either MS2-*β-globin*-UTR or MS2-*Draxin*-UTR (10 min intervals). MS2-containing transcripts are visualized by MCP fluorescence, pseudocolored using a lookup table (LUT) to highlight differences in fluorescence intensity (LUT calibration indicated on left). (**D-E**) Representative confocal maximum intensity projection micrograph of MS2-*Draxin*-UTR transcripts, indicated by MCP fluorescence, colocalized with the fluorescently-tagged P-body marker DCP1a (FP-DCP1a) in an explanted neural crest cell. (**F**) Mander’s coefficient (M1) was calculated to determine the extent of colocalization; 90° rotation of MCP images were compared to unrotated FP-DCP1a images as a control (M1_90_). (**G-H**) Representative epifluorescence images of explanted neural crest cells expressing MS2-*Draxin*-UTR transcripts, indicated by MCP fluorescence, co-electroporated with Cas9 and either control gRNA or *DDX6* gRNA constructs (indicated by H2B-RFP fluorescence). (**I**) Intensity profiles were calculated as normalized gray values across the cytoplasm. Dotted arrows in (**G-H**) indicate example intensity profile measurements. HH, Hamburger-Hamilton stage; MCP, MS2 coat protein; M, Mander’s coefficient; UTR, untranslated region. Scale bars, 1 μm. Error bars, SEM. *, *P* < 0.02, Kolmogorov-Smirnov test. See also **Supplemental Figures S1-S3**.

To determine if the *Draxin* 3’-UTR localizes to P-bodies during cranial neural crest EMT, we first asked whether known P-body components are expressed in these cells. Using HCR *in situ* hybridization, we observed enrichment of P-body-associated transcripts (*e*.*g*., *DCP1a, XRN1, DDX6/ p54*) (Decker and Parker, 2012) in premigratory and migratory cranial neural crest cells as well as neural tissues (**Supplementary Fig. S2**). Given that neural crest cells have the necessary components to form P-bodies, we next asked whether the MS2-*Draxin*-UTR transcripts colocalized with any P-body proteins in migrating crest cells. As commercial antibodies for these proteins failed to work with chick tissue, we co-electroporated a construct expressing human DCP1a fused to the far-red fluorescent protein FP635 (FP-DCP1a) with YFP-tagged MCP and MS2-*Draxin*-UTR and analyzed fixed cranial neural crest explants; the results showed that MS2-*Draxin*-UTR transcripts significantly associated with FP-DCP1a puncta (**Fig. 2D-F**; *P* < 0.001, Kolmogorov-Smirnov nonparametric test), confirming the results inferred by MCP localization.

To further validate the localization of MS2-*Draxin*-UTR transcripts to P-bodies, we used CRISPR/ Cas9 to knockdown the RNA helicase DDX6 (**Supplementary Fig. S3**; 69.4 ± 3.0% of the control, *P* < 0.001, two-tailed paired *t*-test), which has been shown to be required for P-body formation (Andrei et al., 2005; Standart and Weil, 2018). Importantly, *DDX6* knock-down drastically altered the subcellular localization of MS2-*Draxin*-UTR transcripts (**Fig. 2G-I**; *P* = 0.018, Kolmogorov-Smirnov nonparametric test). Together, these data suggest that a DDX6-dependent mechanism recruits *Draxin* transcripts via its 3’-UTR to P-bodies during EMT, and these are likely sites of RNA decay in cranial neural crest.

### RNA decays within P-bodies during neural crest EMT

Use of the MS2/MCP reporter system in neural crest explants allowed us to infer that a critical regulator of EMT—*Draxin*—localizes to P-bodies and is subsequently degraded (**Fig. 2**). However, recent data from cell lines has suggested that P-bodies may act as sites of storage rather than decay; based on the absence of degradation intermediates within P-bodies, as assayed using either fluorescence activated particle sorting (FAPS) with RNA-sequencing (Hubstenberger et al., 2017) or a two-color MS2/PP7 stem loop-based reporter system (TREAT) (Horvathova et al., 2017), previous investigators hypothesized that degradation within these immortalized cell lines occurs within the cytosol rather than in P-bodies. To distinguish between these possibilities in the context of neural crest EMT, we adapted the two-color TREAT reporter assay to the neural crest system to determine whether *Draxin* transcripts are degraded within P-bodies or the cytosol.

To this end, we modified our MS2-*Draxin*-UTR construct that utilized the recently improved MS2 stem loop (Tutucci et al., 2018) (**Fig. 2**) to include the Xrn1-resistant pseudoknots (xrRNA) and PP7 stem loops from the TREAT reporter downstream (3’) of the MS2 loop region (MS2-xrRNA-PP7-*Draxin*-UTR). In this way, Xrn1-mediated decay should occur in a 5’→3’ direction, degrading the MS2/ MCP-GFP fluorescent signal, but will be blocked by the xrRNA pseudoknots, leaving the PP7/PCP-mCherry signal intact. Therefore, sites of decay should accumulate singly fluorescent PP7/PCP-mCherry degradation intermediates. As a control, we used a construct lacking the xrRNA pseudoknots (MS2-PP7-*Draxin*-UTR), which should not accumulate singly fluorescent degradation intermediates (**Fig. 3A**). This follows the original logic of the TREAT reporter system and helps determine to what extent (if any) the PP7/PCP-mCherry complex may artificially stabilize transcripts.

**Figure 3.**
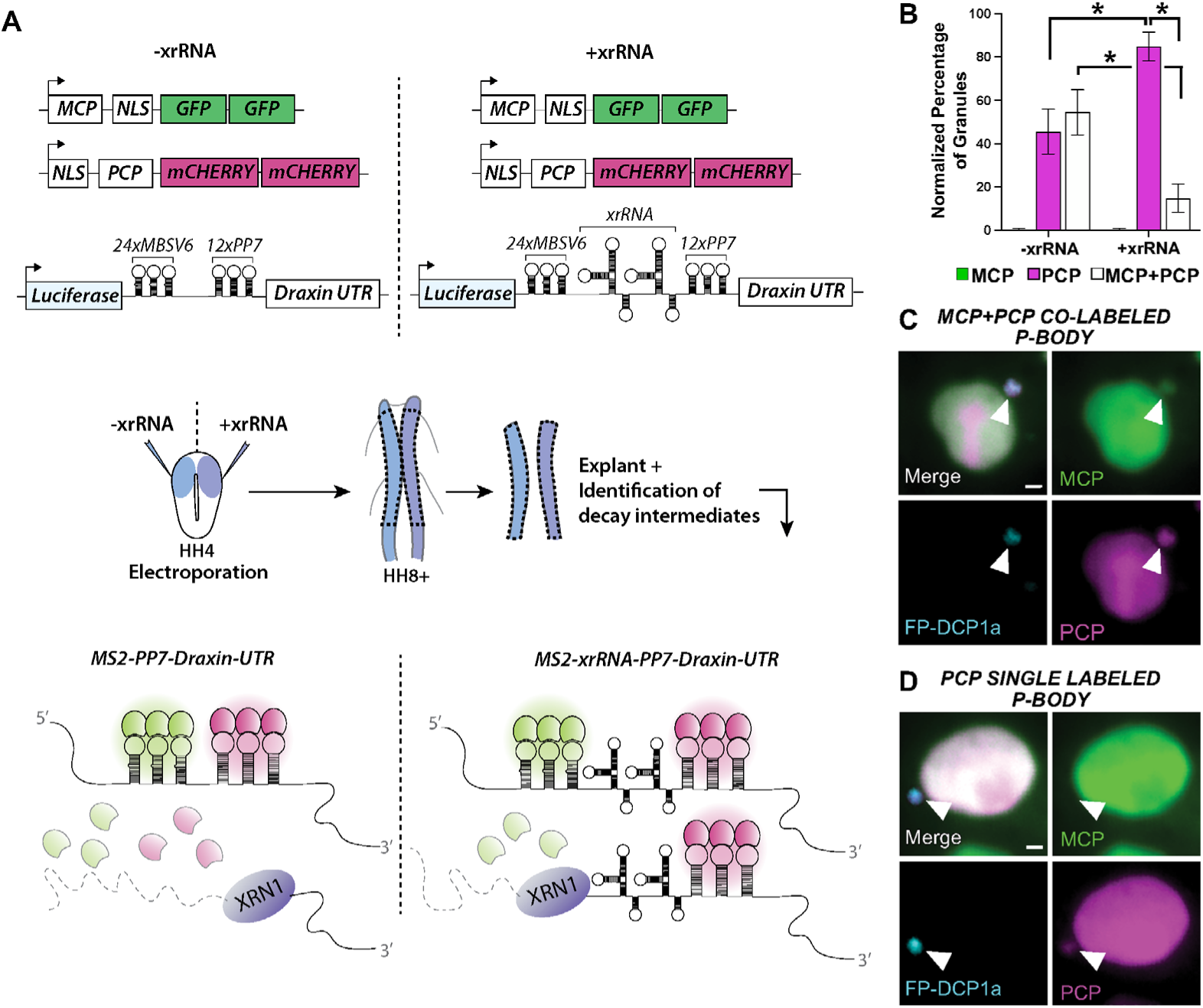
Targeted *Draxin* decay occurs within P-bodies. (**A**) Schematic diagram of expected fluorescence results using a modified TREAT reporter containing the *Draxin* 3’UTR to localize Xrn1-mediated transcript decay. (**B**) Normalized percentage of P-bodies containing singly fluorescent MCP granules, singly fluorescent PCP granules, or dual-fluorescent MCP/PCP granules, for neural crest cells expressing the modified TREAT reporter with/without the xrRNA pseudoknot. (**C-D**) Representative epifluorescence images of dual labeled (MCP+PCP) or single labeled (PCP only) P-bodies in neural crest cells expressing the modified TREAT reporter. MCP, MS2 coat protein; PCP, PP7 coat protein; UTR, untranslated region. Scale bars, 1 μm. Error bars, SEM. *, *P* ≤ 0.01, two-tailed *t*-test. See also **Supplemental Figure S4**.

The results show that the pseudoknot-containing construct (MS2-xrRNA-PP7-*Draxin*-UTR) yields significant enrichment of singly fluorescent PP7/PCP-mCherry degradation intermediates overlapping with the P-body marker (FP-DCP1a) (**Fig. 3B-D**; *P* < 0.001, two-tailed *t*-test, *n* = 128 P-bodies), an indication of 5’→3’ decay. Furthermore, using time lapse live imaging, we observed degradation of the MS2/MCP-GFP fluorescent signal while the PP7/PCP-mCherry signal remained intact within a single P-body (**Supplemental Fig. S4**).

In contrast, the control construct lacking the xrRNA pseudoknot (MS2-PP7-*Draxin*-UTR), failed to yield significant differences (**Fig. 3B**; *P* = 0.51, two-tailed *t*-test, *n* = 60 P-bodies) between singly- or dual-fluorescent MCP/PCP signals within P-bodies, suggesting that the PP7 stem loops in complex with PCP-mCherry may partially inhibit Xrn1-mediated decay, at least in the neural crest. These results suggest that use and positioning of PP7 stem loops in reporter constructs to assess decay in relation to P-bodies should be carefully considered based on context. Importantly however, we observed significant differences for the singly fluorescent PP7/PCP-mCherry (*P* = 0.01, two-tailed *t*-test), as well as the dual-fluorescent MCP/PCP signals (*P* < 0.01, two-tailed *t*-test), with the presence of the xrRNA pseudoknot compared to its absence (**Fig. 3B**). Thus, while the PP7 stem loops may minimally interfere with transcript decay in our system, positioning these loops downstream of the MS2 and xrRNA loops still allowed detection of 5’→3’ decay. From these data, we propose that Xrn1-mediated decay of *Draxin* occurs within P-bodies in cranial neural crest.

### P-body disruption blocks cranial neural crest EMT *in vivo*

Use of an artificial reporter system in neural crest explants allowed us to infer that degradation of a critical regulator of EMT—*Draxin*—occurs within P-bodies (**Fig. 3**), and that P-body localization of the *Draxin* 3’-UTR requires DDX6 (**Fig. 2**). We next asked what happens to endogenous *Draxin* in intact chick embryos when *DDX6* is knocked down. Using HCR, we observed perdurance of endogenous *Draxin* (**Fig. 4A-C**; 113.8 ± 2.5% of the control side, *P* < 0.001, two-tailed paired *t*-test, *n* = 8 embryos), which is normally downregulated during neural crest EMT (**Fig. 1**). Importantly, we found that neural crest EMT was inhibited by loss of *DDX6* (**Fig. 4D**; 86.4 ± 2.7% of the control side, *P* < 0.001, two-tailed paired *t*-test, *n* = 15 embryos), which phenocopies the effects of aberrantly maintaining *Draxin* expression during EMT beyond when it should have been endogenously downregulated (Hutchins and Bronner, 2018). This is consistent with observations in human patients with *DDX6* mutations, who display defects in P-body assembly and often present with craniofacial abnormalities (Balak et al., 2019), a hallmark of cranial neural crest dysfunction (Vega-Lopez et al., 2018). Together, our data highlight a critical role for *DDX6* in the control of cranial neural crest EMT and provide evidence for Xrn1-mediated decay within P-bodies of a biologically relevant target with an important developmental function.

**Figure 4.**
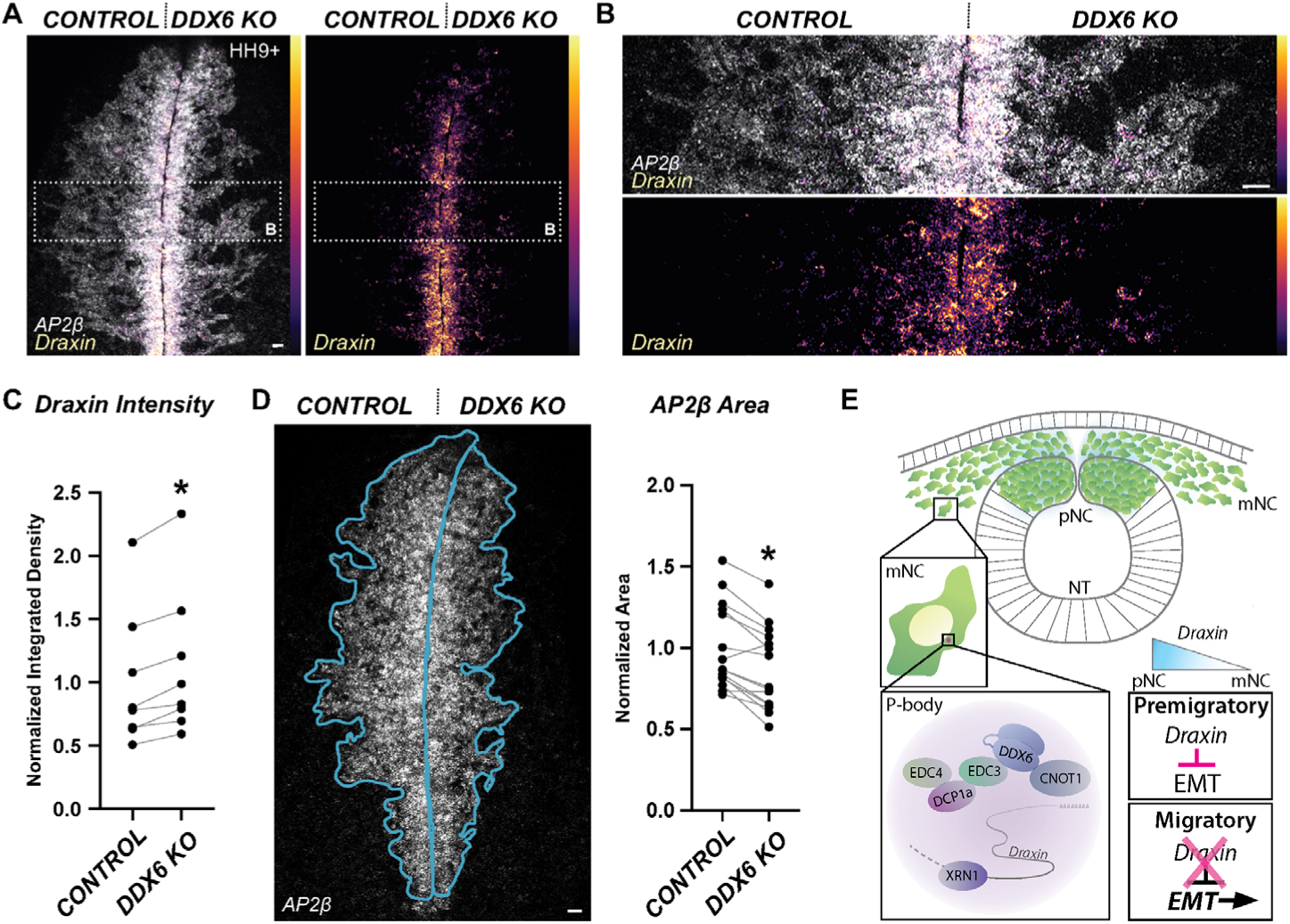
P-bodies are integral to *Draxin* decay and neural crest EMT. (**A-B**) Representative confocal maximum intensity projection micrographs of HCR processed embryos for endogenous *Draxin* and *AP2β* transcripts in whole mount embryos bilaterally electroporated with constructs encoding Cas9 and control gRNA (left) or *DDX6* gRNA (right). *Draxin* transcripts are pseudocolored using a lookup table (LUT) to highlight differences in fluorescence intensity (LUT calibration indicated on right). (**C**) Relative fluorescence intensity of *Draxin* for *DDX6* knockdown compared to control sides of individual embryos. (**D**) Representative confocal maximum intensity projection micrographs of HCR processed embryos for *AP2β* transcripts in whole mount embryos with *DDX6* knockdown, used to calculate the area of neural crest migration (blue outline). Normalized area measurements compared the *AP2β* expression domain for *DDX6* knockdown compared to control sides of individual embryos. (**E**) Model depicting the role of P-bodies in mediating *Draxin* decay to control neural crest EMT. HH, Hamburger-Hamilton stage; KO, CRISPR/Cas9 knockout; pNC, premigratory neural crest; mNC, migratory neural crest; NT, neural tube. Scale bars, 20 μm. *, *P* < 0.001, paired *t*-test.

## DISCUSSION

There is a growing appreciation for the role of post-transcriptional regulation in the control of EMT, particularly in the neural crest. Much of our understanding of the control of EMT has come through careful dissection of gene regulatory networks and transcription-dependent cellular changes during development and disease (Aiello and Kang, 2019; Aiello et al., 2018; Nieto et al., 2016; Yang et al., 2020). Recent studies have suggested that the transcriptional networks controlling EMT are titrated post-transcriptionally via microRNA-mediated repression (Bhattacharya et al., 2018; Cursons et al., 2018; Sanchez-Vasquez et al., 2019; Weiner, 2018). Translational repression or transcript degradation via microRNAs effect subtle adjustments on network gene expression (Bartel, 2018). In stark contrast, we demonstrate here that full and rapid transcript clearance of a molecular rheostat is essential for controlling EMT. Our work further provides the first demonstration in metazoans that this targeted mRNA decay occurs within P-bodies during EMT (**Fig. 4E**).

But why invoke multiple modes of post-transcriptional repression to control EMT? The fine-tuning of expression levels by microRNAs is important in light of recent paradigm shifts in our understanding of EMT—that cells often display partial EMT phenotypes that are highly dynamic, existing on a sliding scale between “epithelial” or “mesenchymal” states (Nieto et al., 2016; Yang et al., 2020). How does this fit with full-scale RNA clearance within P-bodies? In the neural crest, *Draxin* expression acts like a throttle for EMT; once that throttle is removed via P-body-mediated decay, and EMT initiates, we hypothesize a shift to microRNA-mediated repression to influence the spectrum of EMT cell states required for the neural crest to acquire more mesenchymal characteristics and initiate migration. During development, one can imagine similar scenarios directing cell fate choices—whereby dynamic and reversible repression by microRNAs may drive cell fate specification, but cell fate commitment may require transcript clearance by P-bodies.

For microRNA-independent transcript decay, RNA-binding proteins (RBPs) likely mediate transcript localization to P-bodies in neural crest. RBPs recognize *cis* elements within RNA transcripts, particularly in 3’-UTRs, to direct mRNA subcellular localization, translational regulation, or decay (Gebauer et al., 2012). In addition, RBP function and localization is often controlled through post-translational modifications as a result of upstream signaling pathways (Schoenberg and Maquat, 2012). Given that many developmental processes, including neural crest specification and EMT, are downstream of signaling events, RBPs offer a mode of coordination of extra-cellular signals with transcript regulation leading to cell state changes. Understanding the balance of transcript stability versus decay, and the mechanisms mediating these choices, will yield valuable insight into the control EMT, as well as cell fate decisions, during development and disease.

## ACKNOWLEDGMENTS

We thank A. Collazo and G. Spigolon for imaging assistance at the Caltech Biological Imaging Facility; M. Schwarzkopf and G. Shin (Molecular Technologies) for HCR probe design; S. Manohar, G. da Silva Pescador, and C.J. Andrews for cloning assistance; R. Galton for pilot HCR experiments; and R. Singer, J. Chao, and E. Izaurralde for essential reagents made available via Addgene. This work was supported by the National Institutes of Health [R01DE027538 and R01DE027568 to M.E.B; K99DE028592 to E.J.H; K99DE029240 to M.L.P.].

## AUTHOR CONTRIBUTIONS

Project was conceived by E.J.H. and M.E.B. Experimental design and data interpretation were conducted by E.J.H, M.L.P, and M.E.B. Time-lapse experiments were performed by E.J.H. and M.L.P. Chick electroporations and explants were performed by E.J.H. Constructs were designed and generated by E.J.H. and M.L.P. Embryology, hybridization chain reaction experiments, imaging, and quantitation and statistical analyses were performed by E.J.H. Manuscript was written by E.J.H. and M.E.B., with editing by M.L.P.

## METHODS

### Model organism and embryo collection

Fertilized chicken eggs (*Gallus gallus*) were obtained commercially (Sunstate Ranch, Sylmar, CA). Eggs were incubated at 37°C to reach specified Hamburger-Hamilton (HH) stage (Hamburger and Hamilton, 1951). Embryos were collected from eggs using Whatman filter paper as described (Hutchins and Bronner, 2018), and processed as indicated below.

### Hybridization chain reaction, immunohistochemistry, and sectioning

Hybridization chain reaction v3.0 (HCR; Molecular Technologies) was performed according to manufacturer’s instructions (Choi et al., 2018) with custom HCR probe sets and Alexa488, Alexa546, and Alexa647 HCR amplifiers. Immunohistochemistry was performed for Pax7 as previously described (Hutchins and Bronner, 2018). Cross-sections were cryosectioned at 20 μm using previously published protocols (Hutchins and Bronner, 2018, 2019).

### Electroporations and neural crest explants

Electroporations were performed on HH4 gastrula stage chicken embryos as described previously (Hutchins and Bronner, 2018, 2019). Embryos electroporated with reagents for CRISPR/Cas9 were incubated at 37°C for 4 hours immediately following electroporation, then the incubator was shut off for 6 hours to allow CRISPR reagents time to function; incubation at 37°C was then resumed for 8-10 hours to reach the desired HH stage. For all other electroporations, embryos were continuously incubated at 37°C immediately following electroporation until the desired HH stage was reached.

Neural crest explants were generated as described (Manohar et al., 2020; Rogers et al., 2013), with minor modification. Briefly, dorsal neural tubes from 5-6 somite stage (HH8+/HH9-) embryos were dissected in Ringer’s solution. Tissue was then transferred to 1% fibronectin-coated chambered slides (μ-Slide 8 Well Glass Bottom, Ibidi #80827, for live imaging; Nunc™ Lab-Tek™ II Chamber Slide with removable wells, Thermo #154534, for fixed imaging), containing DMEM media supplemented with 10% FBS, 2 mM L-glutamine, and 100 U penicillin/0.1 mg/mL streptomycin. Explants were grown 12-18 hours at 37°C/5% CO_2_, then directly imaged (time lapse experiments) or fixed with 4% paraformaldehyde for 20 min at room temperature. For explants treated with cycloheximide (CHX), CHX (Sigma #C4859) was added to media following overnight incubation to a final concentration of 100 μg/mL; explants were incubated at 37°C/5% CO2 for 1 hour, then washed in PBS and fixed.

### Constructs

Constructs were verified by sequencing prior to electroporation. To generate the d2EGFP-*β-globin*-UTR reporter construct, we excised the IRES-H2B-RFP cassette from pCI-H2B-RFP (Betancur et al., 2010), which contained the *β-globin*-UTR downstream of the cassette. We PCR amplified d2EGFP from M38 TOP-dGFP (Addgene #17114), engineering *Xho*I and *Not*I sites using primer sequences 5’-AAACTCGAGGCCACCATGGTGAGCAAGG and 5’-AAAGCGGC-CGCCTACACATTGATCCTAGCAGAAG and cloned the coding region upstream of the *β-globin*-UTR. To generate the d2EGFP-*Draxin*-UTR reporter construct, we excised the *β-globin*-UTR from the d2EG-FP-*β-globin*-UTR reporter construct using *Not*I and *Pvu*II sites. We cloned in place the *Draxin* 3’-UTR, which we PCR amplified from stage HH9 cDNA using primer sequences 5’-ATAGCGGCCGCGGCTACGCTGTTATGCCAAATTC and 5’-ATACAGCTGCTGCCCCATCCTCAGGTG.

To generate MCP-NLS-2xYFP, we PCR amplified MCP from CYC1p-MCP-NLS-2xyeGFP (Tutucci et al., 2018) (Addgene #104394) using primer sequences 5’-GAATTGCTCGAGGCCACCATGGCTTCTA-ACTTTACTCAGTTCG and 5’-CTCCGGCATCTAC-CCAAAAAAAAAAAGAAAAGTTATCGATATGG, and cloned into the multiple cloning site of pCI-H2B-RFP using engineered *Xho*I and *Cla*I sites. We then excised the IRES-H2B-RFP cassette and cloned in place a 5’ copy of YFP lacking a stop codon, which we PCR amplified from IRES-H2B-YFP-DD (Han et al., 2014) (Addgene #96893) using primer sequences 5’-AAGTTATCGATATGGTGAGCAAGGGCGAGG and 5’-GCATGGACGAGCTGTACAAGAGCGGC-CGCATGGT. We then PCR amplified a 3’ copy of YFP with a stop codon using primer sequences 5’-CAAGAGCGGCCGCATGGTGAGCAAGGG CGAGG and 5’-GGACGAGCTGTACAAGTAAGCGGCCGCAAT-TC and cloned downstream of the first YFP. To generate MCP-NLS-2xGFP, we PCR amplified 2xEGFP from phage UbiC NLS HA stdPCP stdGFP (Horvathova et al., 2017) (Addgene #104099) using primer sequences 5’-TCTATCGATATGGTGAGCAAGGG-CGAG and 5’-ATAGCGGCCGCTTATTTGTACAAT-TCATCCATACCATGGG, and cloned in place of 2xYFP. To generate PCP-NLS-2xmCherry, we PCR amplified PCP from phage UbiC NLS HA stdPCP stdGFP (Horvathova et al., 2017) (Addgene #104099) using primer sequences 5’-ATACTCGAGCGCCAC-CATGGGCCCAAA and 5’-CTCATCGATGGTGGC-GACCGGTGGAC, and cloned into MCP-NLS-2xYFP in place of MCP. We then excised 2xmCherry from pDZ585 pKAN 2x-mCherry (Saroufim et al., 2015) (Addgene # 72236) and blunt cloned the cassette in frame in place of 2xYFP.

To generate the MS2 stem loop constructs, we first PCR amplified the coding region of mTurq2 from pME Turqoise2 (Oehlers et al., 2015) (Addgene #135207) using primer sequences 5’-ATG-GTGAGCAAGGGCGAGGAGC and 5’-ATTGCGG-CCGCTTACTTGTACAGCTCG and cloned into the d2EGFP-*β-globin*-UTR and d2EGFP-*Draxin*-UTR reporter constructs in place of d2EGFP. We next excised the 24xMBSV6 stem loops from pET264-pUC 24xMS2V6 Loxp KANr Loxp (Tutucci et al., 2018) (Addgene #104393) and cloned upstream of the *β-globin*- and *Draxin*-UTRs using *Cla*I and *Not*I sites. To generate the modified TREAT reporter constructs, we first PCR amplified humanized Renilla luciferase from the original TREAT plasmid (Horvathova et al., 2017) (Addgene #104096) using primer sequences 5’-ATACTCGAGCGACTCACTATAGGCTAGCCAC and 5’-ATAGGCCGGCCTTACTGCTCGTTCTTCAG-CAC, and cloned in place of mTurq2 in the MS2-*Draxin*-UTR construct. We then inserted artificial sequence containing unique restriction sites between the 24xMBSV6 stem loops and *Draxin* 3’-UTR to facilitate cloning. We excised the PP7 stem loops from the original TREAT plasmid and blunt cloned between the 24xMBSV6 stem loops and *Draxin* 3’-UTR to generate the Luc-MS2-PP7-*Draxin*-UTR control construct. We then PCR amplified the xrRNA pseudoknots from the original TREAT plasmid using primer sequences 5’-CGCATCGATGCGTA-AGTCAGGCCGGAAA and 5’-GTGACGCGTGTAG-GTAGGATCCTCACCCAGTCC, and cloned into the Luc-MS2-PP7-*Draxin*-UTR control construct between the 24xMBSV6 stem loops and the PP7 stem loops, to generate the Luc-MS2-xrRNA-PP7-*Draxin*-UTR construct.

To generate the FP-DCP1a construct, we PCR amplified the coding region of the far-red fluorescent protein turboFP635 (Evrogen) using primer sequences 5’-TGTACCGGTCTCGAGGCCACCAT-GGTGGGTGAGG and 5’-AGATCCGGAGCTGTG-CCCCAGTTTGCTA, and cloned in place of EGFP in pT7-EGFP-C1-HsDCP1a (Tritschler et al., 2009) (Addgene # 25030). We then excised the FP635-DCP1a cassette and blunt cloned into d2EGFP*β-glo-bin*-UTR in place of the d2EGFP coding region.

For CRISPR/Cas9 experiments, plasmids encoding Cas9, Cas9-GFP, and control gRNA were described previously (Gandhi and Bronner, 2018; Gandhi et al., 2017). The *DDX6* gRNA plasmid was generated by cloning a protospacer sequence (5’-AGATC-GAGAATTCTTCCAG), designed using CHOPCHOP (Labun et al., 2019), into the modified gRNA backbone (Gandhi et al., 2017). For electroporations with untagged Cas9, pCI-H2B-RFP was co-electroporated to infer which cells received the knockdown reagents, as described previously (Gandhi et al., 2017).

### Image acquisition and analysis

Confocal images were acquired using an upright Zeiss LSM 880, an inverted Zeiss LSM 800, and/or an inverted Zeiss LSM 710 at the Caltech Biological Imaging Facility, and epifluorescence images were acquired using a Zeiss Imager.M2 with an ApoTome.2 module. Time lapse experiments were performed using the Zeiss LSM 800 and/or the Zeiss LSM 710 with incubation set to 37°C/5% CO_2_. Images were minimally processed for brightness/contrast and pseudocolored using Fiji (ImageJ, NIH) and Adobe Photoshop 2020.

Colocalization measurements were determined by calculation of the Mander’s coefficients M1 and M2 using the JaCoP (Bolte and Cordelieres, 2006) plugin in Fiji. Mander’s coefficients were calculated using square ROIs approximately 100 μm^2^. For P-body M1 measurements, ROIs were cropped around P-bodies to exclude nuclei. Only cells containing both FP-DCP1a-positive granules and MCP-positive nuclei were considered; within these cells, all P-bodies were measured for colocalization with cytoplasmic MCP signal. As a control for P-body colocalization, ROIs from MCP images were rotated 90° and compared to unrotated FP-DCP1a images to calculate the Mander’s coefficient (M1_90_) to rule out nonspecific overlap by chance (Lee et al., 2020).

Relative area of neural crest migration was determined in Fiji. For each whole mount image, the line tool was used to draw an ROI surrounding the area of neural crest indicated by *AP2β* expression (example indicated in **Fig. 4D**). Measurements were made for the control electroporated (left) and experimental electroporated (right) sides from the same embryo, and then the experimental electroporated area was divided by the control electroporated area to calculate the relative area of neural crest migration. Relative fluorescence intensity was determined similarly following background subtraction (50-pixel rolling ball radius), using the same ROIs from area measurements. Relative fluorescence intensity was then calculated by dividing the integrated density measurements for the experimental versus the control side of the same embryo. For time lapse experiments, integrated density was determined for single granules, then normalized per granule over time by dividing measurements by maximum measured integrated density. Normalized integrated density was then averaged across multiple granules per time point for each condition.

### Statistical analysis

Statistical analyses were performed using Prism (8; GraphPad Software). *P* values are defined in the text, and significance was established with *P* < 0.05. For assays determining whether samples have same statistical distribution (**Fig. 2**), significance was assessed using a Kolmogorov-Smirnov nonparametric test. For all other assays of multiple comparisons, *P* values were calculated using two-tailed unpaired (**Fig. 3**) or paired (**Fig. 4**) *t*-tests. Data shown as bar graphs are presented as mean values, with error bars indicating SEM and individual data points shown. Data comparing relative values of control versus morphant sides are plotted as normalized individual values with lines connecting values measured from the same embryo. Number of embryos/samples and replicates are indicated in figure legends and/or text. Data were assumed to be normally distributed but were not formally tested.

## Data availability

All data generated and/or analyzed during the current study are available from the corresponding author on reasonable request.

## Code availability

Information for accessing the JaCoP plugin used to calculate Mander’s coefficients is available: https://imagej.nih.gov/ij/plugins/track/jacop.html

---

**Supplementary Figure S1.**
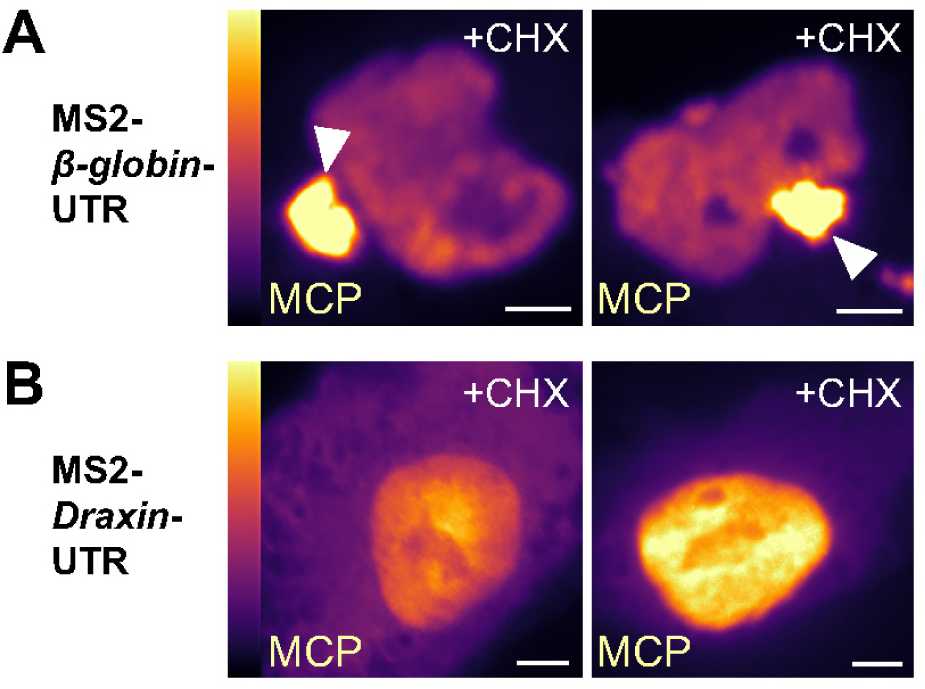
Cycloheximide treatment disrupted granule formation of *Draxin*-UTR-containing transcripts, but not *β-globin*-UTR-containing transcripts. (**A**) Representative epifluorescence images of explanted neural crest cells expressing MS2-*β-globin*-UTR and YFP-tagged, nuclear MCP, treated with 100 μg/mL CHX. (**B**) Representative epifluorescence images of explanted neural crest cells expressing MS2-*Draxin*-UTR and YFP-tagged, nuclear MCP, treated with 100 μg/mL CHX. MS2-containing transcripts are visualized by MCP fluorescence, pseudocolored using a lookup table (LUT) to highlight differences in fluorescence intensity (LUT calibration indicated on left). Arrowhead, presence of cytoplasmic granule. MCP, MS2 coat protein; CHX, cycloheximide. Scale bars, 5 μm.

**Supplementary Figure S2.**
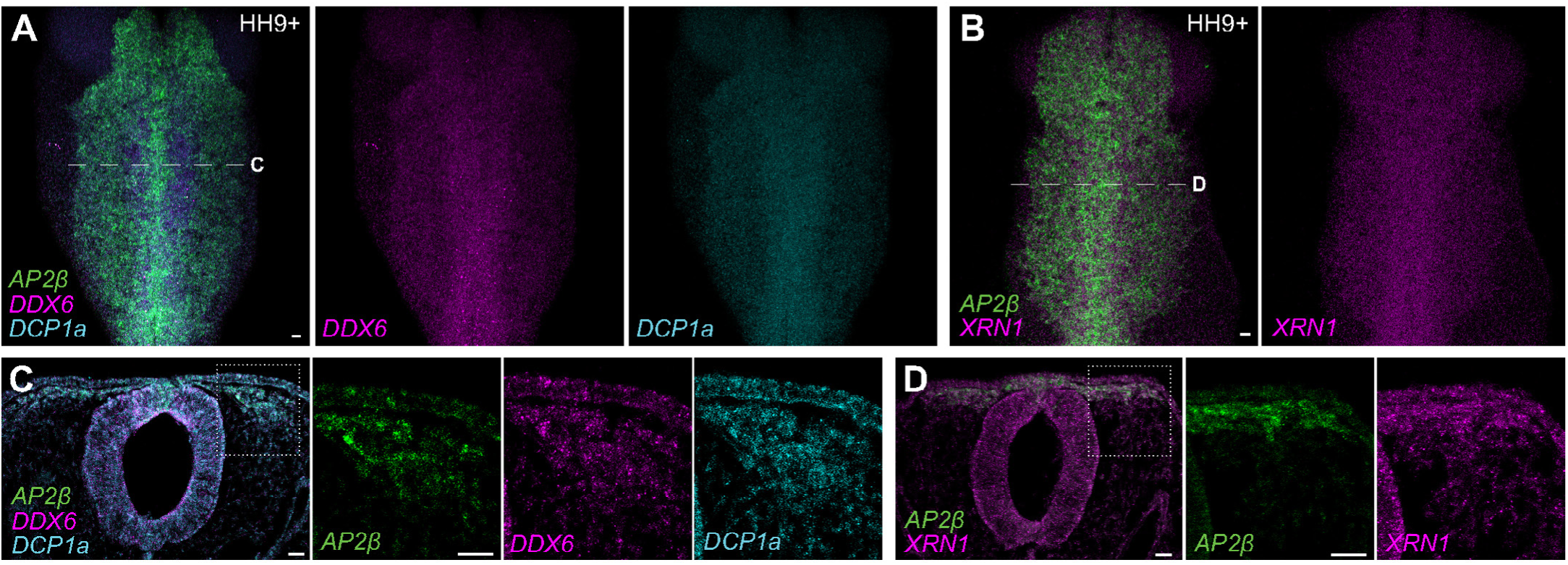
P-body components are enriched in migratory neural crest cells. (**A-D**) Representative confocal maximum intensity projection micrographs of HCR processed wild type embryos for P-body components and neural crest marker *AP2β* transcripts in whole mount (**A-B**) and cross-section (**C-D**). Boxed areas in (**C-D**) indicate zoomed regions in right panels. HH, Hamburger-Hamilton stage. Scale bars, 20 μm.

**Supplementary Figure S3.**
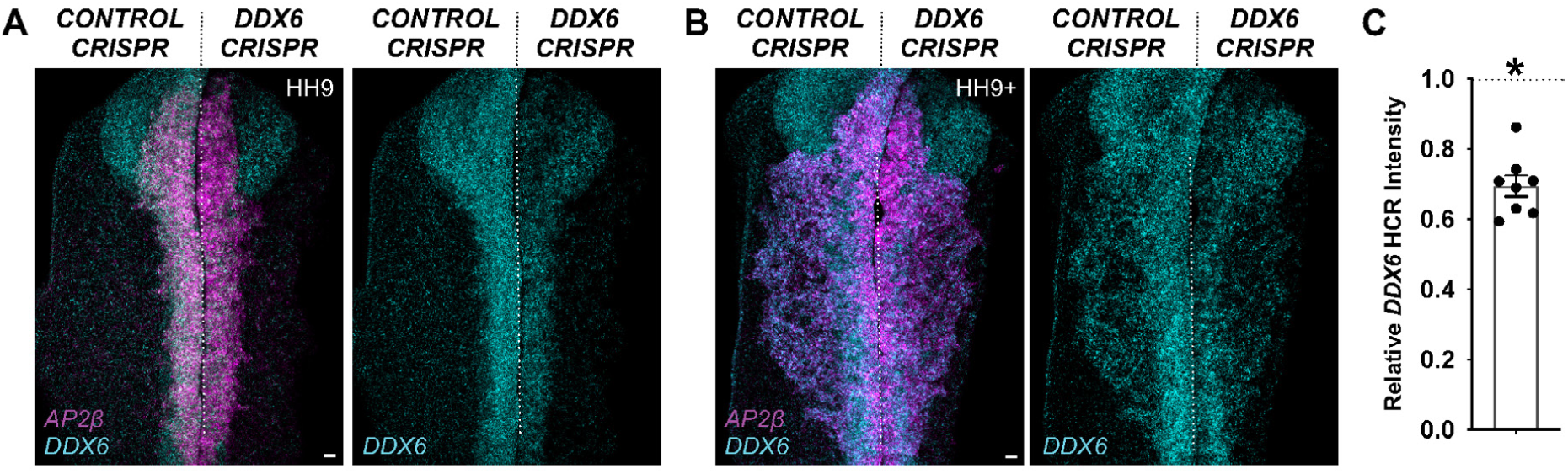
Validation of CRISPR/Cas9-mediated knockdown of *DDX6*. (**A-B**) Representative confocal maximum intensity projection micrographs of HCR processed embryos for *DDX6* and neural crest marker *AP2β* transcripts in whole mount embryos bilaterally electroporated with constructs encoding Cas9 and control gRNA (left) or *DDX6* gRNA (right). (**C**) Mean relative fluorescence intensity of *DDX6*, calculated as the ratio of normalized integrated density in *DDX6* knockdown versus control side for individual embryos. HH, Hamburger-Hamilton stage. Scale bars, 20 μm. Error bars, SEM. *, *P* < 0.001, two-tailed paired *t*-test.

**Supplementary Figure S4.**
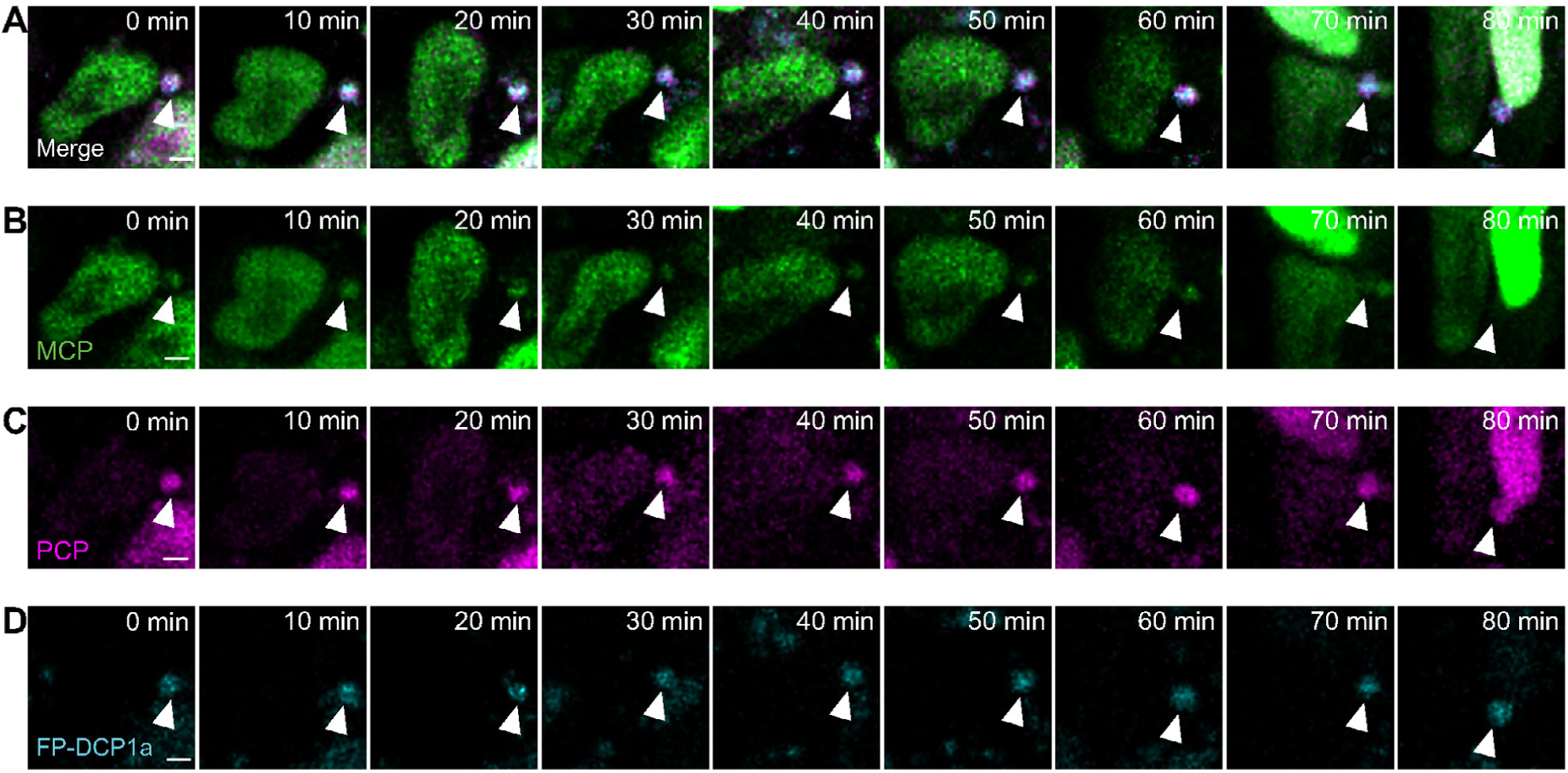
Live RNA imaging of Draxin decay within a P-body in a migratory neural crest cell. (**A-D**) Time lapse confocal micrographs of decay intermediates of MS2-xrRNA-PP7-*Draxin*-UTR transcripts, indicated by MCP/PCP fluorescence, colocalized with the fluorescently-tagged P-body marker DCP1a (FP-DCP1a) in an explanted neural crest cell. MCP, MS2 coat protein; PCP, PP7 coat protein. Scale bars, 1 μm. Error bars, SEM. *, *P* < 0.001, two-tailed paired *t*-test.

